# One organ, two engines: divergent mitochondrial phenotypes in anterior and posterior crab gills

**DOI:** 10.64898/2026.05.25.727634

**Authors:** Georgina A. Rivera-Ingraham, Mariel Familiar-López, Gillian Renshaw

**Affiliations:** Australian Rivers Institute. Griffith University. 4215. Edmund Rice Drive. 4215. Southport, Queensland Australia; UMR 9190-MARBEC. Centre National de la Recherche Scientifique (CNRS). Place Eugène Bataillon. 34090. Montpellier, France; School of Environment and Sciences, Griffith University. 4215. Parklands Drive, Southport, Queensland Australia; Graduate School, University of Queensland. 4072. St Lucia, Queensland, Australia; School of Pharmacy and Medical Science, Griffith University. 4215. Parklands Drive. Southport, Queensland, Australia

## Abstract

Gills are multifunctional organs that integrate respiration with homeostasis, including energy demanding processes such as osmoregulation and excretion. In osmoregulating decapod crustaceans, two spatially segregated gill types differ in function, ultrastructure and membrane composition, as well as in their responses to environmental change. Yet mitochondrial function in crab gills remains poorly characterized.

Here, for the first time, we used high-resolution respirometry with a substrate–uncoupler–inhibitor titration (SUIT) protocol to characterize the mitochondrial phenotypes in anterior (respiratory) and posterior (osmoregulatory) gills. For this, gill filaments of the shore crab *Carcinus maenas* were permeabilized for 30 min in a saponin solution (optimized for each tissue at 25 µg or 5 µg saponin · mg^−1^ gill fresh weight for anterior and posterior gills, respectively).

Anterior gills exhibited higher leak control ratios (L/P, L/E), consistent with a leak-dominated mitochondrial phenotype that may contribute to redox balance at expense of maximal ATP yield. In contrast, posterior gills, displayed a higher phosphorylation control ratio and tighter coupling (higher Net P), reflecting a tightly-coupled, ATP-producing mitochondrial phenotype, in line with their role in sustaining ATP-intensive activities such as osmoregulation and excretion.

Our results revealed that anterior and posterior gills operate as “two engines in one organ”: by quantifying how each gill type partitions respiratory capacity between phosphorylation and leak pathways, this study provides a mechanistic framework for understanding how mitochondrial specialization supports functional division of labour within a single organ and contributes to physiological adaptation to dynamically fluctuating marine environments.

## INTRODUCTION

Gills are multifunctional, highly metabolic organs that integrate respiration with essential homeostatic processes, encompassing ion and osmoregulation, ammonia excretion, and acid–base balance (Evans et al., 2005; Henry et al., 2012; Wood et al., 1995). To fulfill this broad spectrum of functions, crustaceans possess two spatially segregated gill types: anterior gills primarily support gas exchange and contribute to ammonia excretion (Weihrauch et al., 1998), whereas posterior gills are specialized for ion uptake (e.g. Siebers et al., 1982) and osmoregulation (Copeland and Fitzjarrell, 1968; Gilles et al., 1988; Péqueux, 1995). This functional division is accompanied by marked differences in ultrastructure, membrane composition, and physiological responses to environmental stressors.

Beyond structural and functional specialization at the tissue level, anterior and posterior gills also differ in their redox physiology. For instance, under exposure to diluted seawater, posterior gills significantly upregulate antioxidant defenses and prevent reactive oxygen species (ROS)-induced cellular damage, whereas anterior gills seemingly lack such response and show increased susceptibility to oxidative damage (Rivera-Ingraham et al., 2016). These differences suggest that each tissue may operate under distinct constraints in terms of energy metabolism and redox balance, raising the possibility that mitochondrial function itself is differentially tuned between gill types.

Mitochondria play a central role in integrating energy production with redox homeostasis through the control of electron transport, proton leak and ATP synthesis. In many systems, mitochondrial function is not only tissue-specific but also reflects the balance between ATP demand, ROS production and metabolic flexibility (Fernández-Vizarra *et al*., 2011), and that such differences can become even more pronounced under pathological conditions (Balmaceda *et al*., 2024). By extension, one may expect similar tissue-specific patterns in estuarine organisms, which are regularly exposed to dynamic, severe environmental fluctuations. However, despite the well-established functional divergence between anterior and posterior gills, the mitochondrial mechanisms underpinning this specialization remain largely unexplored. Understanding these differences would provide valuable insight into why certain tissues, such as anterior gills, are more vulnerable to environmental stressors.

Here, we hypothesized that mitochondrial function differs systematically between anterior and posterior gills, reflecting their contrasting physiological roles. Specifically, we predicted that: i) anterior gills would display higher relative proton leak and lower coupling efficiency, indicative of a greater allocation of electron flux to redox balance rather than ATP production; ii) posterior gills would exhibit higher oxidative phosphorylation capacity and tighter mitochondrial coupling, consistent with the sustained ATP demand for ion transport; and iii) these differences would be evident at the level of mitochondrial control ratios, reflecting distinct allocation of electron transport system (ETS) capacity rather than differences in mitochondrial abundance.

To test these hypotheses, we used high-resolution respirometry and a substrate-uncoupler-inhibitor titration (SUIT) protocol to characterize mitochondrial respiratory states, flux control ratios and coupling efficiency in permeabilized anterior and posterior gills, using the European shore crab *Carcinus maenas* as a study model. By quantifying how each gill type partitions respiratory capacity between phosphorylation and leak pathways, this study provides a mechanistic framework for understanding the functional specialization within a single organ and its relevance to environmental adaptation in marine invertebrates.

## MATERIALS AND METHODS

### Animal collection and maintenance

Specimens of *C. maenas* (L.) were collected from Victoria (Southeastern Australia) using round crab pots consisting in four entrance funnels made of heavy duty 15mm nylon mesh and with an overall measurement of 800mm wide x 300 mm heigh. These pots were baited with sardines and were deployed from the shoreline and allowed to sit in the seabed for a maximum of 2h. Only male crabs were used in this study to avoid interference of reproduction-related energetic differences between sexes. Crabs were transported to Griffith University’s laboratory facilities at Gold Coast, Queensland. Animals were maintained under in 150-L seawater (SW) aquaria equipped with a water pump, aeration bubblers and a filtration pump (1200 L/h) in a controlled temperature room at 15ºC. Animals were fed thawed fish, twice weekly. Every 48 h, water quality was checked using Quantofix® nitrate/nitrite test strips (Macherey-Nagel GmbH and Co., Duren, Germany). Routinely, a partial water change was carried out weekly unless nitrate and/or nitrite values increased over 10 and 1 mg l^−1^, respectively. Within aquaria, animals were maintained in 920ml individual perforated BPA-free plastic boxes 15 × 15 × 5 cm (l × w × h) to avoid social conflicts among crabs.

### Experimental setup and tissue sampling

Animals were fed and immediately transferred to smaller aquaria (8L), where they remained isolated and free from their boxes for 7d. Water in these smaller aquaria was maintained at normoxia (>99% air saturation) via air bubblers. The water was changed daily.

After 7 days, the animals (N=7, average carapace width 6.4 ± 0.2cm; average weight 63.1 ± 7.6g) were rendered insensitive to stimuli by placing them in a freezer for 10 min. Crabs were euthanized using a spike through the brain.

### Haemolymph osmolality

For each of the crabs analysed, a small haemolymph sample (~200 µl) was obtained using a 1ml hypodermic needle inserted at the base of the last pereiopod. Samples were immediately used to assess the osmotic pressure of the haemolymph through freezing point depression osmometry using a Osmomat 3000basic (Gonotec, Berlin, Germany). In addition, hemolymph osmolality was used as a standard to ensure that all buffers used for the high-performance respirometry were isosmotic.

### Gill sampling and preparation

This study used a paired design, where both anterior gills (pairs 1-4, representative of purely respiratory gills) and posterior gills (pairs 7-8, as representative of osmoregulatory gills) were dissected from each crab and placed in a modified ice-cold relaxing buffer (2.77 mM CaK_2_EGTA, 7.23 mM K_2_EGTA, 5.77 mM Na_2_ATP, 6.56 mM MgCl_2_·6H_2_O, 20 mM taurine, 20 mM imidazole, 0.5 mM dithiothreitol, 50 mM MES, and 400 mM KCl, pH 7.1). This buffer, originally designed for mammalian tissues, was modified by omitting phosphocreatine and adding approximately 400 mM KCl to increase the osmolarity to match the osmolality of crab hemolymph (≈ 920 mOsm · kg^−1^) following the method of Iftikar et al. (2010).

Ice cold gill samples were blotted dry, weighed and mechanically chopped using fine scissors. Following optimized protocols for saponin dose and incubation time determined by Rivera-Ingraham et al (submitted), anterior and posterior gills were permeabilized by incubating tissues in ice-cold relaxing buffer containing either 25 µg saponin · mg gill FW^−1^ or 5 µg saponin · mg tissue FW^−1^ respectively. The incubation time for both anterior and posterior gill tissues was 30 min over ice, on an orbital shaker at 80 rpm. Then, samples were washed three times in respiration buffer (0.5 mM EGTA, 3 mM MgCl_2_, 100 mM K-lactobionate, 20 mM taurine, 10 mM KH_2_PO_4_, 20 mM HEPES, 200 mM sucrose, 250 mM KCl and 1 g L^− 1^ BSA, pH 7.24 at 20 °C, 995 mOsm) for 10 minutes over ice, on an orbital shaker. All chemicals were purchased from Sigma Aldrich.

### Assessment of mitochondrial function

Mitochondrial function was determined for anterior and posterior gills separately, using individual chambers in an Oxygraph-2K (Oroboros Instruments, Innsbruck, Austria). Chambers were held at 15ºC with continuous gentle stirring. Initially, we ran a series of parallel studies to determine the preferred mitochondrial complex I (CI) substrates for crabs. The substrates tested differed not only in chemical nature but also their electron entry routes, the following three substrates were tested: proline (a proteinogenic aminoacid) (10 mM) and malate (a TCA cycle intermediate) (5 mM), for providing good flux rates in other crustaceans and other marine invertebrates (Iftikar et al., 2010; Liu et al., 2013; Steffen et al., 2023), as well as pyruvate (a glycolytic end product) (5 mM). Since the highest CI contribution to oxidative phosphorylation (OXPHOS) was obtained with proline, it was retained for this study. Because the addition of pyruvate with malate often failed to induce an increase in oxygen flux, yet malate had a slight impact on OXPHOS. Consequently, proline and malate were included in all SUIT protocols, as in Iftikar et al. (2010).

Gills, with an average weight 17.8 ± 1.8 mg were allowed to sit in the dark with gentle stirring for 20 minutes to allow the consumption of endogenous substates. Then the first reading of oxygen flux was taken to establish the baseline for routine respiration (R_baseline_). A substrate–uncoupler–inhibitor-titration (SUIT) protocol was commenced to measure the respiration rates of mitochondrial complexes, electron leak states, as well as the isolated contribution of mitochondrial complexes CI and CII to OXPHOS, illustrated in Figure 1, the subsequent calculations from the SUIT protocol are presented in table 1.

**Table 1:**
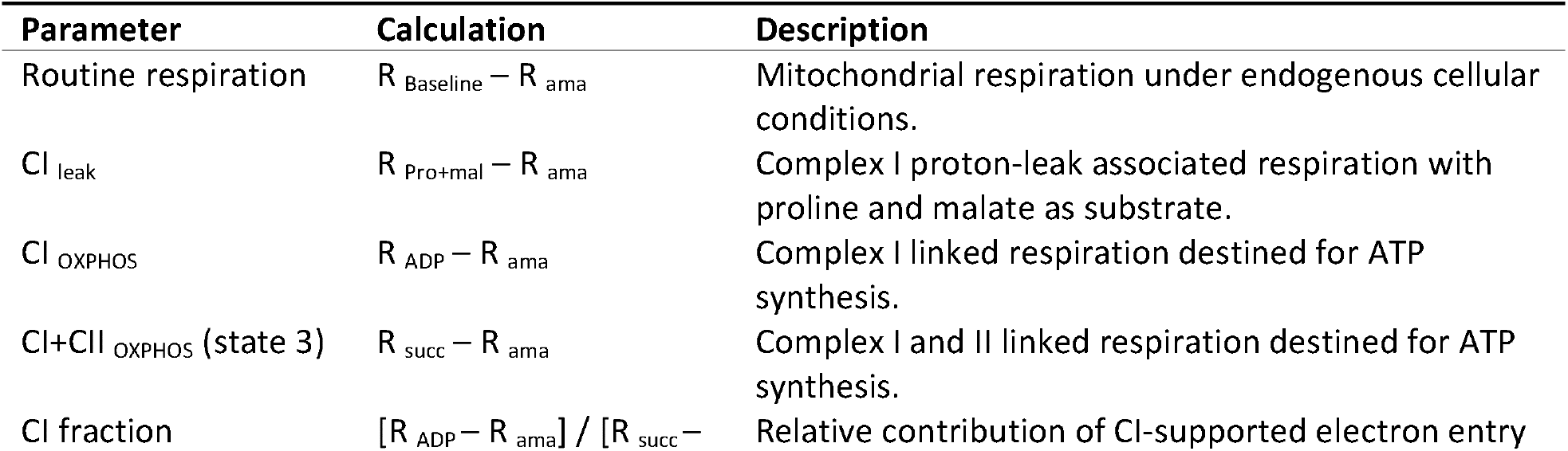
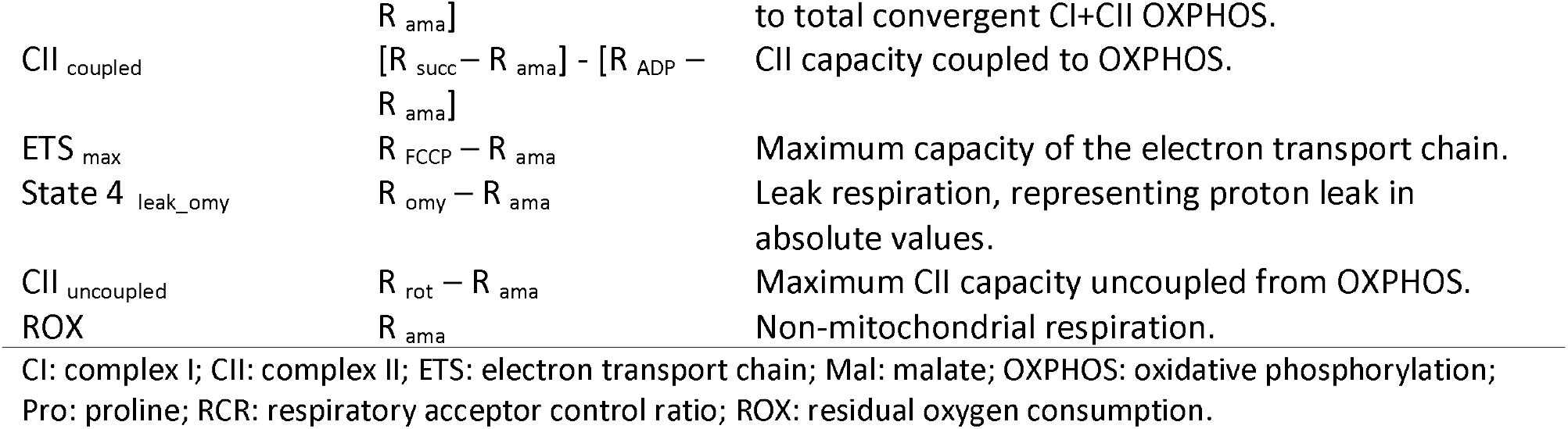
Absolute mitochondrial respiratory fluxes (corrected fluxes for non-mitochondrial oxygen consumption) and pathway-specific capacity descriptors measured in permeabilized anterior and posterior gills of Carcinus maenas.

**Figure 1:**
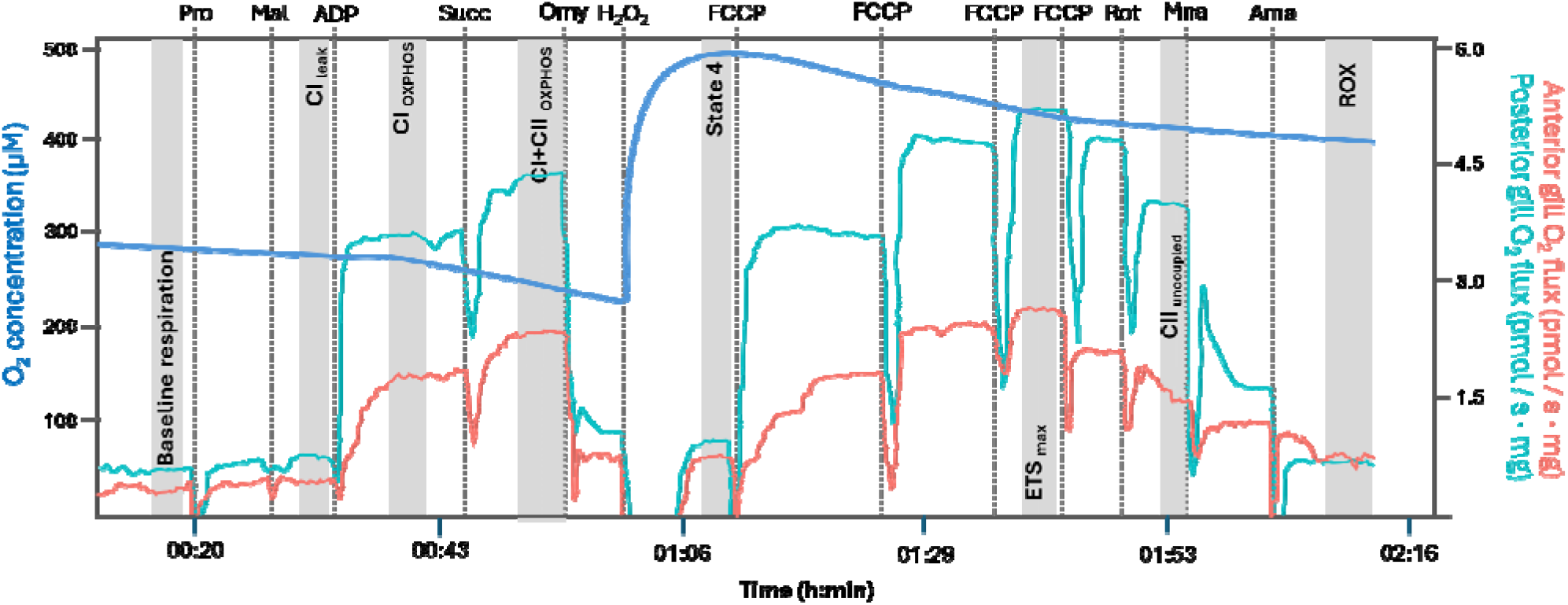
Representative trace of a mitochondrial respiration assay of permeabilized gill fibers from C. maenas. The dark blue line represents the O_2_ concentration while the red and light blue lines show the mitochondrial respiratory flux normalized to tissue mass for anterior and posterior gills, respectively. The parameters measured in this study are written vertically in the body of the graph. Substrates and inhibitors used are marked at the top, with a dashed line indicating moment of titration. Ama: antimycin; CAT: catalase; CI: mitochondrial complex I; CII: mitochondrial complex II; FCCP: carbonylcyanide-p- (trifluoromethyl)phenylhydrazone; Mal: malate; Mna: malonic acid; Omy: oligomycin; Pro: proline; Rot: rotenone; ROX: residual (non-mitochondrial) oxygen consumption; S: sample; Suc: succinate.

In the first step, CI substrates proline (10 mM) and malate (5 mM) were added to each chamber before the addition of ADP, to measure CI, state 2 respiration (CI _leak_) (Figure 1, Table 1). ADP was added in excess (1 mM) to measure the CI contribution to OXPHOS (CI _OXPHOS_). Then the CII substrate succinate was added at a concentration of 10 mM to assess the convergent electron flow during maximal OXPHOS respiration (CI+CII _OXPHOS_ or State 3). The addition of Oligomycin (Omy) (6 µM) was used to inhibit ATP synthase (complex V) and determine the mitochondrial respiration associated with ATP synthesis (State 4). The maximal flux of the electron transport system (ETS _max_) was determined by uncoupling mitochondrial respiration by using a multiple-step titration of carbonylcyanid-p- (trifluoromethyl)phenylhydrazone (FCCP) at a concentration of 0.2 µM. FCCP was added until respiration flux decreased with regards of the previous titration. Then the CI inhibitor rotenone (Rot) (1.25 µM), was titrated to determine the isolated capacity of CII to fuel the ETS with electrons (CII _uncoupled_) (Gnaiger, 2020). Finally, the inhibitors malonate (Mna) (15 mM) and antimycin-a (Ama)(6 µM) were titrated to achieve a shutdown of CII and CIII activities, respectively, to determine the non-mitochondrial residual oxygen consumption (ROX).

In order to isolate functional differences from mitochondrial abundance, anterior and posterior gill fluxes were ROX-corrected and expressed as flux-control ratios (FCRs) relative to ETS capacity (calculated as in Doerrier and Gnaiger, 2016) (Table 2).

**Table 2:** Flux control ratios (FCRs) and coupling efficiency indices in permeabilized anterior and posterior gills of Carcinus maenas.

| FCR | Calculation | Description |
| --- | --- | --- |
| CII/E | $[R_{\text{rot}} - R_{\text{ama}}] / [R_{\text{FCCP}} - R_{\text{ama}}]$ | Indicator of how dominant the CII pathway is relative to the ETS<sub>max</sub>. |
| CII/P | $[R_{\text{rot}} - R_{\text{ama}}] / [R_{\text{succ}} - R_{\text{ama}}]$ | Indicator of how much phosphorylating respiration could be, in principle, sustained by CII alone. |
| L/E | $[R_{\text{omy}} - R_{\text{ama}}] / [R_{\text{FCCP}} - R_{\text{ama}}]$ | Leak FCR |
| L/P=1/RCR | $[R_{\text{omy}} - R_{\text{ama}}] / [R_{\text{succ}} - R_{\text{ama}}]$ | L/P Coupling-control ratio (Gnaiger, 2020), representing the fraction of OXPHOS capacity that is lost to proton leak when ATP synthase is inhibited. |
| P/E | $[R_{\text{succ}} - R_{\text{ama}}] / [R_{\text{FCCP}} - R_{\text{ama}}]$ | Phosphorylation system control ratio (Gnaiger, 2020), representing the degree to which OXPHOS is limited by the phosphorylation system. |
| R/E | $[R_{\text{Baseline}} - R_{\text{ama}}] / [R_{\text{FCCP}} - R_{\text{ama}}]$ | Routine FCR, indicating how close routine respiration operates to ETS<sub>max</sub>. |
| Net P | $\begin{aligned} &= (P-L)/P \\ &= 1-(L/P) \\ &= ([R_{\text{succ}} - R_{\text{ama}}] - [R_{\text{omy}} - R_{\text{ama}}]) / [R_{\text{succ}} - R_{\text{ama}}] \end{aligned}$ | OXPHOS coupling efficiency, ranging between 0.0 (zero coupling) to 1.0 (fully coupled system) (Gnaiger, 2020). |

### Determination of mitochondrial density

Mitochondrial density was measured by quantifying citrate synthase (CS) activity in each sample of permeabilized anterior or posterior gills used for each crab, following the method of Srere et al. (1963). Aliquots of each gill sample used were homogenized 1:10 (w:v) in MiRO5, using an Omni Internation Tx homogeniser operated for 5 cycles of 5 sec each at 4ºC. The CS activity was measured in duplicates in 6.5 μl of supernatant and using a final volume of 165 μl. Baseline activity (in the absence of oxaloacetic acid) and free Co-A production (generated following the addition of oxaloacetic acid) were read for each sample as changes in absorbance at 412 nm for at least 3 min at 20°C using a Perkin Elmer EnSight multimode microplate reader (Perkin Elmer, MA, USA). Results were expressed as units (U, μmol min^−1^) per mg of tissue fresh weight.

### Data analysis

Data acquisition and analysis of mitochondrial function was carried out using Oroboros DatLab (Oroboros Instruments, version 7.4.0.4). Data are reported as mean ± standard error of the mean.

Because both anterior and posterior gills from each crab were used in the study, the resulting measurements were not independent within animals. After ensuring that all datasets complied with the assumptions for parametric statistics (Shapiro and Levene tests for normality and homoscedasticity of data), gill type was analyzed for non-mitochondrial oxygen consumption (ROX) using paired t-test (or paired Wilcoxon signed-rank test for non parameteric datasets).

Finally, multivariate patterns in mitochondrial phenotype were explored using principal component analysis (PCA). Variables included in the analysis were selected a priori to represent complementary dimensions of mitochondrial function, including respiratory capacity (e.g. CI-OXPHOS, ETS max), coupling efficiency (Net P), proton leak allocation (L/E), and reserve respiratory capacity (R/E), while minimizing redundancy among highly correlated parameters. Prior to analysis, variables were standardized (z-score transformation) to account for differences in scale and variance among respiratory parameters. PCA was performed using the prcomp function in R, and scores from the first two principal components were used to visualize tissue-specific clustering of anterior and posterior gill mitochondrial phenotypes. Variable loadings were projected onto ordination space to facilitate interpretation of the major axes of mitochondrial functional variation. Differences among phenotypes in multivariate space were tested using permutational multivariate analysis of variance (PERMANOVA; adonis2 R package, Euclidean distance, 999 permutations).

All statistical analyses and figure representations were conducted in R (v.4.4.0) using Rstudio “Cucumberleaf Sunflower” Release (Posit Software, PBC).

## RESULTS

Mitochondrial density, assessed via CS activity did not differ significantly between anterior and posterior gills (t = −1.01, *p* = 0.357) (Fig. 2), indicating that subsequent differences in respiratory parameters reflect functional variation rather than differences in mitochondrial abundance.

**Figure 2:**
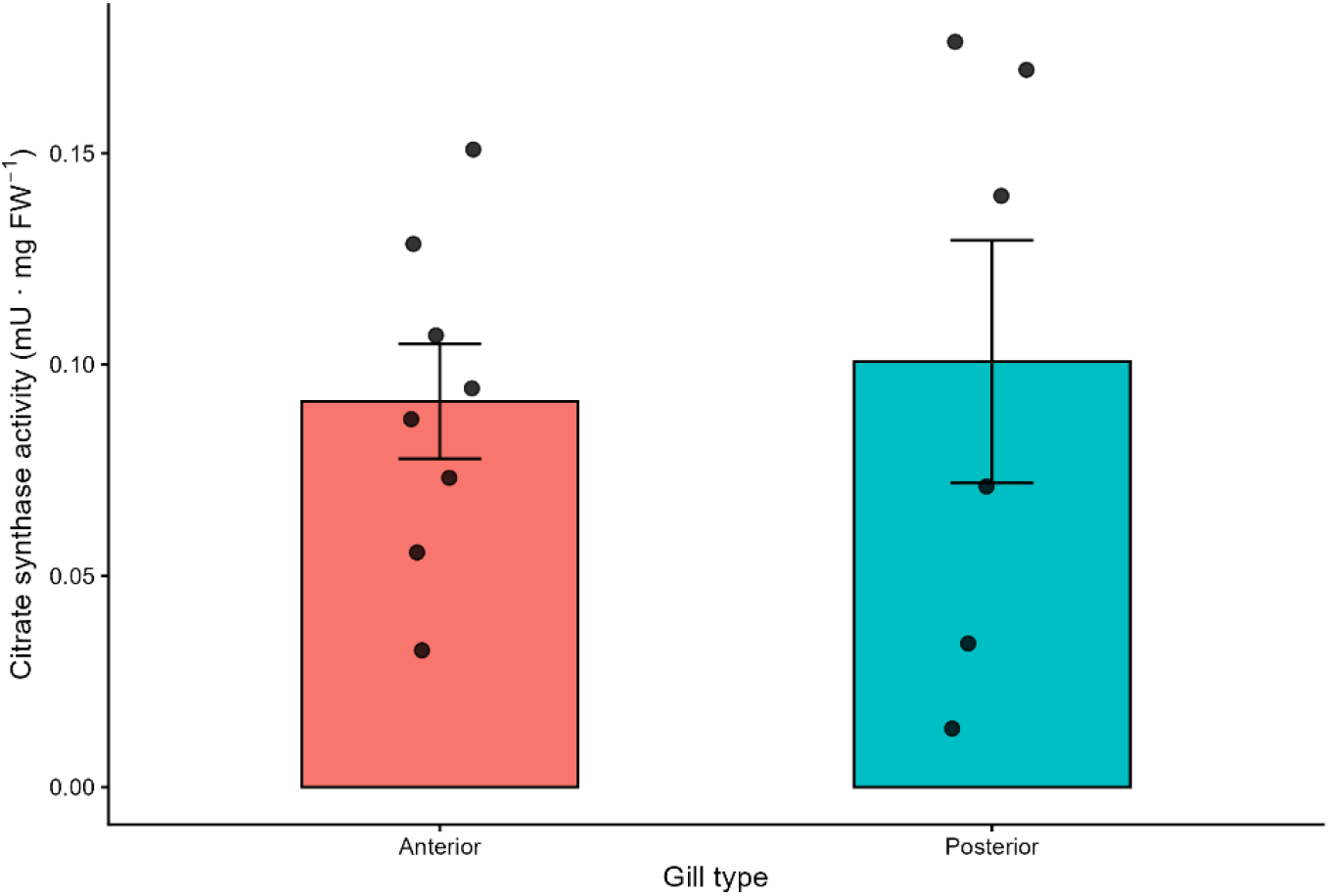
Citrate synthase activity measured in anterior and posterior gills of each of the crabs considered in the study. Bars represent the standard error of mean. Black dots depict individual data points. Statistical analyses show no significant differences between anterior and posterior gills (t = −1.01, *p* = 0.357).

### Tissue-specific differences in mitochondrial fluxes

Clear and consistent differences in mitochondrial respiratory fluxes were observed between anterior and posterior gills (Table 3; Fig. 3). When compared to anterior gills, posterior gills exhibited significantly higher rates of oxidative phosphorylation, including Complex I-linked respiration (CI-OXPHOS; p = 0.005), combined Complex I + II respiration (State 3; p = 0.006), and maximal electron transport system capacity (ETS max; p = 0.006). In addition, posterior gills had significantly higher CII-supported uncoupled respiration (p = 0.022), than anterior gills.

**Table 3:** Summary statistics and paired comparisons of mitochondrial parameters between anterior and posterior gills. Values are presented as mean ± standard error of mean. Differences (Δ) correspond to paired contrasts calculated as posterior-anterior values for each individual. Statistical comparisons were performed using t-paired tests, with the choice of test based on the normality of paired differences. Significant differences are shown in bold.

| | Parameter | Average ± SE | | $\Delta$ (Post-Ant) | Test | df | p-value |
| --- | --- | --- | --- | --- | --- | --- | --- |
|  |  | Anterior gill | Posterior gill |  |  |  |  |
| Net fluxes | Routine | 0.528±0.05<br>7 | 0.683±0.12<br>2 | 0.155 | -1.16679 | 3 | 0.328 |
| | CI-OXPHOS | 1.952±0.27<br>5 | 5.480±0.88<br>5 | 3.528 | -5.67919 | 4 | **0.005** |
| | CI-Leak | 0.872±0.16<br>6 | 1.122±0.12<br>5 | 0.250 | -2.55745 | 4 | 0.063 |
| | ETS max | 3.142±0.46<br>9 | 8.316±1.42<br>0 | 5.174 | -5.2121 | 4 | **0.006** |
| | State 3 | 2.665±0.35<br>8 | 6.876±1.10<br>1 | 4.211 | -5.22323 | 4 | **0.006** |
| | State 4 | 0.840±0.19<br>2 | 1.082±0.18<br>0 | 0.242 | -3.52216 | 4 | **0.024** |
| | CI-fraction | 0.730±0.03<br>8 | 0.796±0.013 | 0.066 | -0.93026 | 4 | 0.405 |
| | CII-uncoupled | 1.438±0.27<br>6 | 4.792±1.08<br>6 | 3.354 | -3.62159 | 4 | **0.022** |
| Ratios | Net P | 0.639±0.05<br>9 | 0.845±0.014 | 0.206 | -4.08414 | 3 | **0.026** |
| | R/E | 0.168±0.03<br>1 | 0.079±0.019 | -0.089 | 3.318391 | 3 | **0.045** |
| | L/P | 0.361±0.05<br>9 | 0.155±0.014 | -0.206 | 4.084139 | 3 | **0.026** |
| | L/E | 0.314±0.05<br>2 | 0.133±0.021 | -0.181 | 3.337396 | 3 | **0.044** |
| | P/E | 0.856±0.03<br>2 | 0.850±0.090 | -0.006 | 0.09632 | 4 | 0.928 |
| | CII/E | 0.442±0.03<br>3 | 0.574±0.056 | 0.132 | -1.48254 | 4 | 0.212 |
| | CII/P | 0.517±0.03<br>5 | 0.699±0.084 | 0.182 | -1.68057 | 4 | 0.168 |

Note: due to occasional assay instability or incomplete traces, sample size varied slightly among parameters

**Figure 3:**
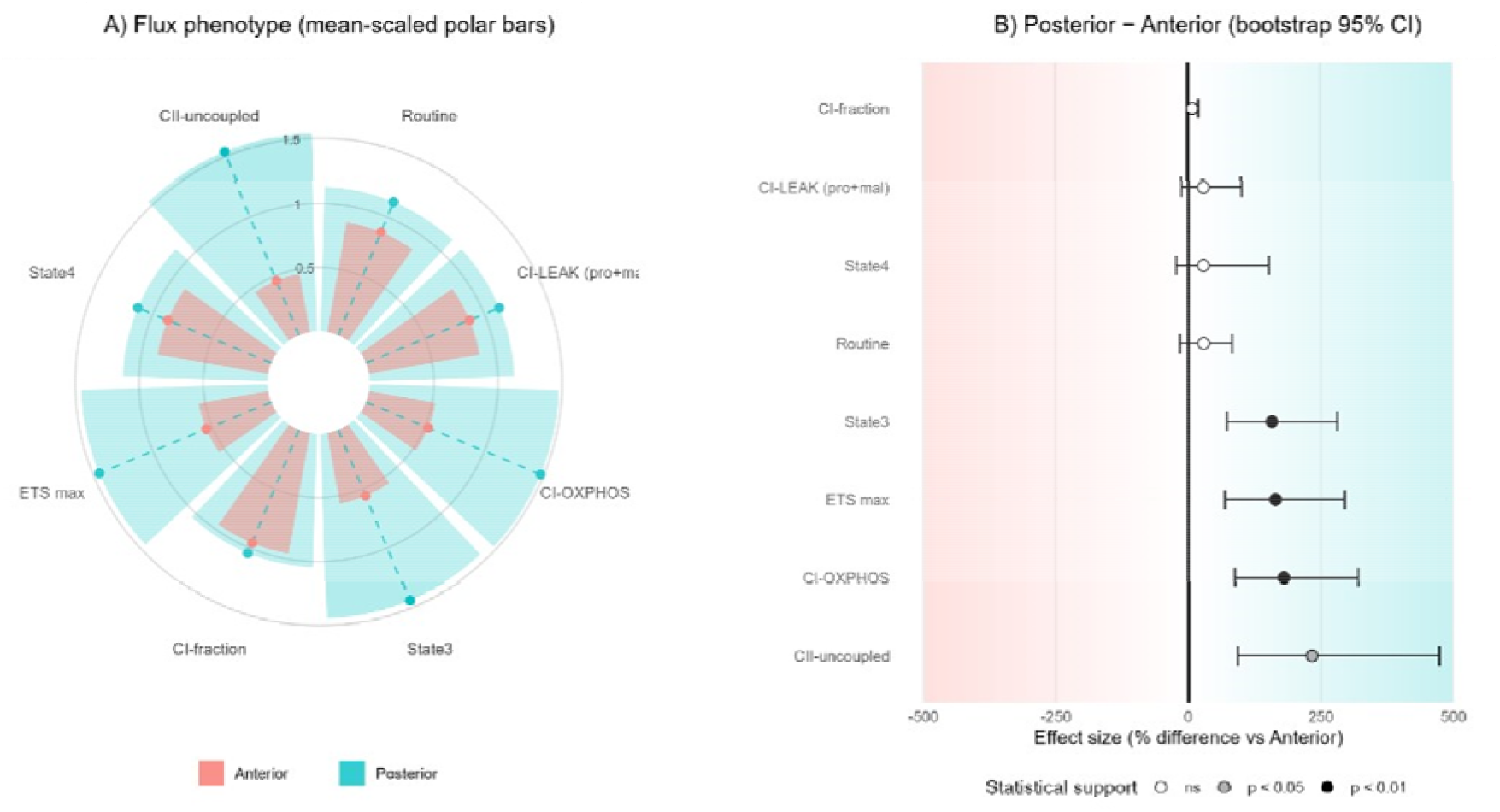
Tissue-specific mitochondrial flux phenotypes in anterior and posterior gills of C. maenas. A) Circular plot showing mean-scaled mitochondrial flux parameters measured in permeabilized anterior (red) and posterior (blue) gills (see Table 2). For each parameter, values are scaled to the mean across tissues (mean =1) to emphasize relative differences in flux allocation between gill types. Posterior gills are shown as wide blue bars, with anterior gills overlaid as narrower red bars. Dashed radial lollipops indicated the magnitude of each tissue-specific mean. B) Effect-size plot showing posterior-anterior differences for each flux parameter, expressed as percentage differences relative to anterior gills. Points indicate mean effect sizes and horizontal lines denote 95% bootstrap confidence intervals. Background color intensity increases with the absolute magnitude of the effect, with direction indicating anterior-biased (left-red) or posterior-biased (right, blue) fluxes. Point gill encodes statistical support from paired tests (see legend) (pair t-tests for all parameters except for State 4, which was analyzed using a paired High on test)

While leak respiration appeared to differ between tissues, only State 4 respiration (oligomycin-induced leak) was significantly higher in posterior compared to anterior gills (*p* = 0.024). CI-linked leak respiration showed a non-significant trend toward higher values in posterior gills (*p* = 0.063), Routine respiration did not differ significantly between tissues (*p* = 0.328), and the relative contribution of Complex I to OXPHOS (CI fraction) remained unchanged (*p* = 0.405).

Overall, these results indicate that, compared to anterior gills, posterior gills operate at higher absolute respiratory capacities across multiple mitochondrial states, while maintaining a similar relative contribution of CI to total OXPHOS.

### Differences in mitochondrial coupling and efficiency

Flux control ratios (FCRs) and coupling indices revealed marked differences in mitochondrial efficiency between tissues (Table 3; Fig. 4). Posterior gills displayed significantly higher OXPHOS coupling efficiency, than anterior gills (Net P; *p* = 0.026), revealing that a greater proportion of respiration was devoted to ATP synthesis. In contrast, anterior gills exhibited significantly higher leak-related ratios, including L/P (*p* = 0.026) and L/E (*p* = 0.044) compared to posterior gills, reflecting a greater relative contribution of proton leak to total respiration.

**Figure 4:**
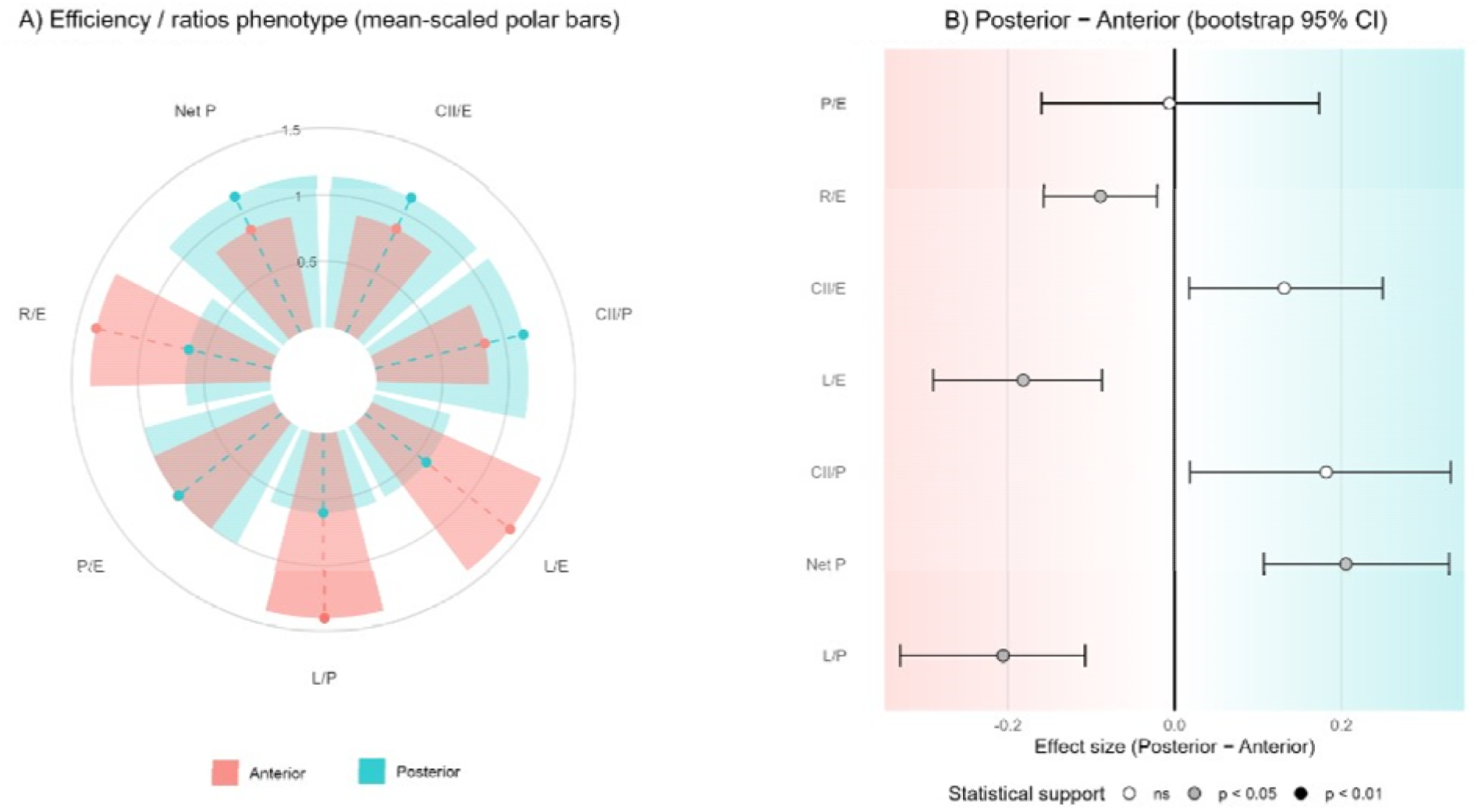
Tissuse-specific mitochondrial efficiency and coupling ratios in anterior and posterior gills. A) Circular plot showing mean-scaled mitochondrial efficiency and coupling ratios measured in permeabilized anterior (red) and posterior (blue) gills (see Table 3). For each parameter, values are scaled to the mean across tissues (mean =1) to emphasize relative differences in flux allocation between gill types. Posterior gills are shown as wide blue bars, with anterior gills overlaid as narrower red bars. Dashed radial lollipops indicated the magnitude of each tissue-specific mean. B) Effect-size plot showing posterior-anterior differences for each flux parameter, expressed as absolute differences. Points indicate mean effect sizes and horizontal lines denote 95% bootstrap confidence intervals. Background color intensity increases with the absolute magnitude of the effect, with direction indicating anterior-biased (left-red) or posterior-biased (right, blue) fluxes. Point gill encodes statistical support from paired tests (see legend) (pair t-tests).

Routine respiration relative to ETS capacity (R/E) was significantly higher in anterior gills (*p* = 0.045), revealing that these tissues operate closer to their maximal respiratory capacity. In contrast, posterior gills had a lower R/E values, indicative of a greater reserve capacity. No significant differences were observed for phosphorylation system control (P/E; *p* = 0.928), nor for the relative contribution of Complex II to ETS capacity (CII/E; *p* = 0.212) or OXPHOS (CII/P; *p* = 0.168).

### Substrate-specific characteristics of CI respiration

Proline was clearly the preferred CI substrate: in anterior gills, addition of proline increased respiration by 8.50 ± 6.38 % over baseline values, while in posterior gills, the increase in respiration was 36.77 ± 11.48 %. Furthermore, these differences were statistically significant (t=-4.915; p<0.01), revealing that proline had a greater effect on the respiration of posterior gills. The addition of malate did not consistently stimulate Complex I-linked respiration in either anterior or posterior gills. In approximately half of the measurements, the addition of malate resulted in a slight but non-significant decrease in oxygen flux, which may suggest a limited or context-dependent contribution of this substrate under the experimental conditions.

The principal component analysis (PCA) identified a strong separation between anterior and posterior gill mitochondrial phenotypes, with PC1 explaining 84.6% of the total variance (Figure 5A). Positive PC1 scores were associated with higher CI OXPHOS, ETS max and NetP, consistent with a tightly coupled and ATP-oriented phenotype. Conversely, anterior gill mitochondria, clustered negative for PC1, and were associated with a higher L/E and R/E, indicative of a lower-coupling phenotype operating closer from their ETS capacity. PC2 explained 8.2% of the total variation, contributing little to overall tissue discrimination. Furthermore, plotting leak allocation relative to ETS capacity (L/E) against routine respiration relative to ETS capacity (R/E) further separated anterior and posterior gills, with anterior gills showing higher leak allocation and operating closer to their ETS capacity, whereas posterior gills showed lower leak allocation and greater apparent reserve capacity (Figure 5B). This tissue-specific separation was further supported by the significant results of a PERMANOVA analysis (pseudo-F=10.78; p<0.01).

**Figure 5:**
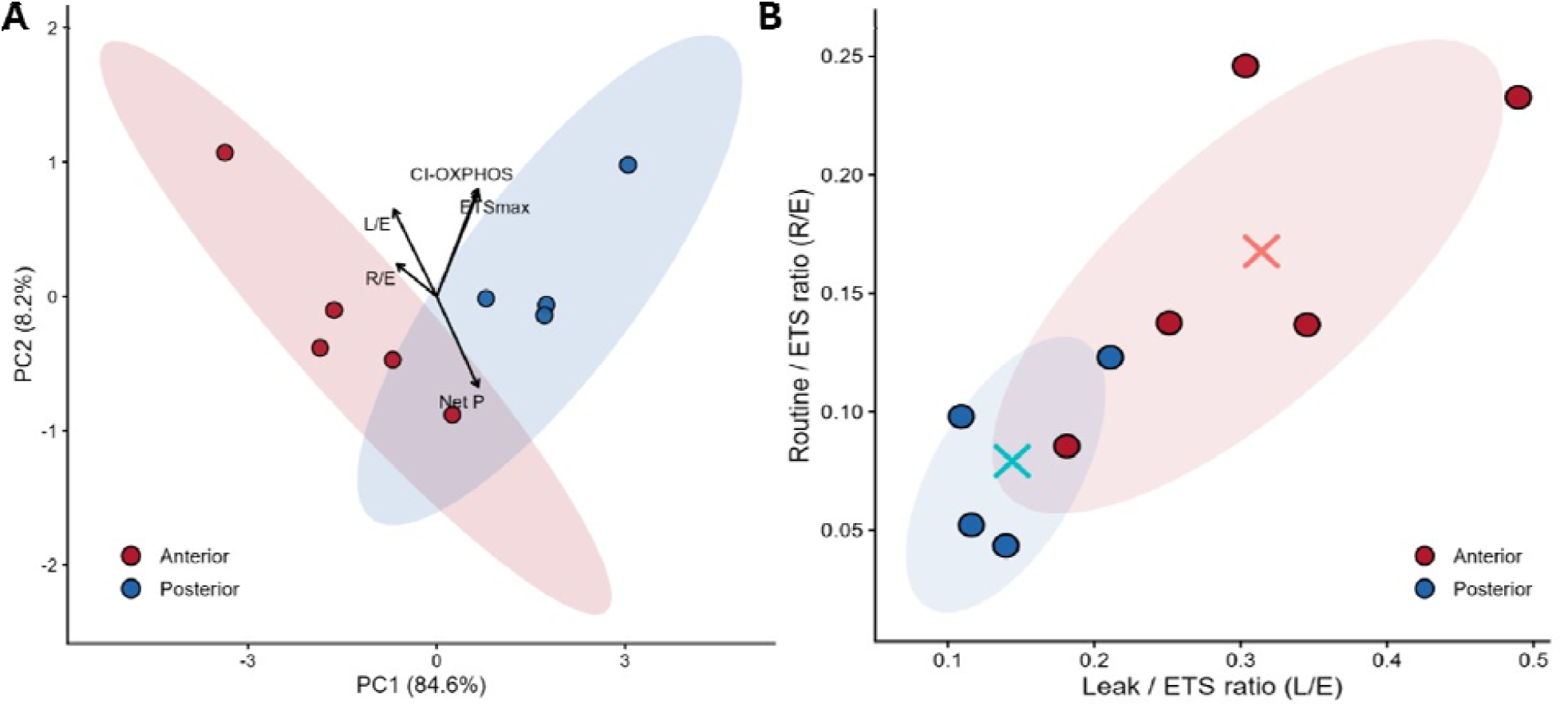
Integrative representation of mitochondrial phenotypes in anterior and posterior gills of C. maenas. A) Principal component analysis (PCA) of mitochondrial phenotype based on variables selected a priori to represent complementary dimensions of mitochondrial function, including respiratory capacity (CI_OXPHOS_, ETS_max_), coupling efficiency (Net P), proton leak allocation (L/E), and reserve respiratory capacity (R/E). Variables were standardized (z-score transformation) to account for differences in scale and variance among respiratory parameters. B) Mitochondrial allocation phenotype represented as the relationship between L/E and R/E. Ellipses represent the multivariate dispersion of each gill type and are shown for visualization purposes only; crosses represent centroids. Each point represents an individual, gill preparation.

## DISCUSSION

### Location-specific division of labour in gill tissue: mitochondrial phenotypes linked to capacity and/or efficiency

Our results provide new insights into the bioenergetic specialization of anterior versus posterior gills. Taken together, the results reveal that anterior and posterior gill mitochondria exhibit functional mitochondrial divergence, independent of mitochondrial density. The flux measurements in conjunction with flux control ratios, show that these two tissues do not just differ in their respiratory capacity, but also in how their electron transport system (ETS) capacity is divided differently between a focus on ATP production (posterior gills) vs a focus on leak-controlled pathways (anterior gills), which lower ROS production. These data support the concept of metabolic specialization within a single organ, whereby anterior gills with higher leak ratios and ion flux appear to be optimized for dissipative respiration and redox stability, while posterior gills had significantly higher rates of oxidative phosphorylation in accord with their role in ATP-demanding ion transport.

### Anterior gills: priorizing low-coupling and redox control over maximal ATP yield

The flux data revealed that, compared to posterior gills, anterior gills had significantly decreased CI-OXPHOS, State 3, ETS max combined with a higher leak ratios (L/E, L/P), a characteristic of diminished respiratory capacity (Doerrier and Gnaiger, 2016). More specifically, their high R/E values indicate that anterior gills operate close to their ETS capacity, suggesting reduced respiratory capacity. More specifically, these patterns not only reveal reduced coupling efficiency but also suggest that mitochondrial function in anterior gills is not optimized for maximal ATP production but rather that they are operating closer to their ETS ceiling. These data support our first hypothesis, that the mitochondria of anterior gills are proton-leak enriched (low coupling), potentially contributing to redox balance.

Proton leak, while lowering ATP synthesis is associated with a reduction in mitochondrial membrane potential and a concomitant decrease in ROS generation, thereby having a cytoprotective role (reviewed by Brookes, 2005). Thus, the function of higher leak observed in anterior gills may have a prime regulatory redox balance to limit oxidative stress, even at the expense of energetic efficiency. This interpretation is consistent with our previous observations, showing stable ROS dynamics across salinity gradients in anterior gills (Rivera-Ingraham et al., 2016). Functionally, this mitochondrial phenotype aligns with the role of anterior gills as respiratory surfaces, where redox homeostasis may be prioritized over maximizing ATP yield. Operating close to the ETS capacity reflects a system tuned for steady throughout rather than energy capacity.

### Posterior gills: channeling respiratory capacity towards ATP synthesis

Our data revealed that mitochondria from posterior gills had: (i) a higher oxidative phosphorylation capacity (increased CI-OXPHOS); (ii) convergent CI+CII OXPHOS (State 3); and (iii) maximal ETS capacity, all suggestive of a greater capacity for electron transport and ATP synthesis in posterior gill mitochondria.

Furthermore, posterior gills displayed lower leak-related flux control ratios (L/E, L/P) implying a larger fraction of ETS capacity channeled to phosphorylation and ATP production at rest rather than dissipated as proton leak. Together with the higher coupling efficiency (Net P), results suggest that posterior gill mitochondria are functionally optimized to sustain ATP-intensive processes, such as ion transport or osmoregulation. Importantly, posterior gills operated further from their ETS maximum (lower R/E), leaving reserve headroom to respond rapidly to acute ATP demands—consistent with the ionoregulatory work of posterior gills during osmotic challenges (Guo et al., 2024; Henry et al., 2012; Rivera-Ingraham et al., 2016). These data support our second hypothesis: that posterior gills have mitochondria optimized for ATP production.

### Functional divergence reflects allocation of ETS capacity rather than mitochondria abundance

Citrate synthase activity, a commonly used indicator of mitochondrial number, was not significantly different between anterior and posterior gills, indicating that there was a comparable abundance of mitochondria in each gill type. Hence by expressing respiratory parameters as FCRs relative to ETS capacity, the observed differences reveal how each tissue partitions electron flux between phosphorylation and leak pathways. Anterior gills allocate a larger fraction to dissipative processes, whereas posterior gills allocate greater fraction of ETS to ATP synthesis. Taken together these data support our third hypothesis that differences in mitochondrial function reflect allocation strategies rather than differences in mitochondrial density. This distinction is critical, as it demonstrates that mitochondrial specialization in gills can be achieved through the functional tuning of bioenergetic pathways rather than simply increasing mitochondrial content. Such allocation strategies likely reflect the contrasting energetic and redox demands imposed by respiration vs ion transport.

### Distinct substrate utilisation patterns in crustacean gill mitochondria

While proline metabolism is perhaps best known for its important role in sustaining insect flight (e.g. Teulier et al., 2016) and locomotory activity (e.g. Gäde and Auerswald, 2002). Our study revealed that CI-supported respiration in C. maenas gill mitochondria relies strongly on the aminoacid proline as a substrate. Proline is well recognized as an alternative metabolic source in several other animal groups (McDonald et al., 2018), and similar substrate preference has previously been shown in crustaceans (Iftikar et al., 2010) and cephalopods (e.g. Hochachka, 1995).

Across ectothermic animals, stress (e.g. hypoxia, desiccation, and some chemical stresses) usually impairs mitochondrial complex function or shift metabolism away from NADH-linked substrates (e.g. Adzigbli et al., 2024; Sokolova, 2023). Hence, a greater reliance on proline oxidation could potentially contribute to physiological resilience under conditions where CI activity becomes constraint. Indeed, proline oxidation can supply electrons downstream of CI, directly at ubiquinone, and potentially sustain ATP generation even when CI function is compromised (Pallag et al., 2022).

An additional observation emerging from this study was the inconsistent effect of malate on CI-supported respiration. In approximately half of the assays, the addition of malate following proline resulted in a slight decrease in oxygen flux rather than the expected stimulation of Complex I-linked respiration. Although exploratory, these findings highlight the importance of considering the functional impact of the substrate mix on CI function in each tissue and condition.

### Integration of mitochondrial subtypes with gill physiology and the osmorespiratory compromise

The two mitochondrial sub-types investigated and characterized in our study map closely onto the well-established functional and structural divergence between anterior and posterior gills. Indeed, posterior gills, while still respiratory tissues, are primary sites of ion exchange, and consequently exhibit an ultramorphological compromise that must satisfy both efficient gas exchange (thin, highly permeable surfaces) and osmoregulatory control (thicker, less permeable domains), in line with the osmorespiratory compromise, a major theme in fish physiology (Wood and Eom, 2021). To achieve this, posterior gills in fish, display multiple, mitochondria-related adaptations, all geared to sustain ATP demanding work essential to fuel the large Na^+^/K^+^-ATPase activities that are key to sustain osmoregulation. It should be noted that, in crustacea, posterior gills have 3.5 time higher activity of than anterior gills, and these levels increase with decreasing environmental salinity (reviewed by Henry et al., 2012). For example, gills have been reported to change mitochondrial density, namely within chloride cells (also known as mitochondria-rich cells), and which are involved in the uptake/intake of ions according to environmental salinity. Concordantly, in crustaceans, the number of mitochondria have been reported to increase in response to energy demanding processes such as decreases in environmental salinity (Guo et al., 2024; Rivera-Ingraham et al., 2016), which would be consistent with an elevated ATP demand for ion pumping. Spatial organization and diffusion geometry have also been reported to change: ultramorphological observations of posterior gills have revealed mitochondrial behaviour to be unique in salt-regulating tissues: when confronted with osmotic stress, epithelial cells form a well-developed complex of basolateral infolds (where Na^+^/K^+^-ATPase resides) and to where mitochondria migrate and cluster in close contact with the epithelial cells infolds of osmoregulatory tissue (Barra et al., 1983; Compere et al., 1989), minimizing ATP/ADP diffusion distances.

Together, our data extend the concept of the osmorespiratory compromise beyond structural and physiological specialization to the level of mitochondrial function. The contrasting coupling and flux allocation patterns observed between anterior and posterior gills suggest that tissue-specific bioenergetic organization contributes directly to supporting the distinct functional demands of respiration and ion regulation in crustacean gills.

## CONCLUSIONS

Biologically, by normalizing ETS, we show that allocation strategy (phosphorylation vs leak at a given ETS ceiling) differs between tissues independently of mitochondrial density. More specifically, anterior and posterior gills appear to be two engines in one organ, responding to a division of labor: one tuned for safe (low-coupling) respiration (higher leak share, lower phosphorylation share), the other tuned for ATP-demand spikes (higher phosphorylation share, lower leak), consistent with the broader osmorespiratory compromise concept (Wood and Eom, 2021) and our previous work on differential stress vulnerability (Rivera-Ingraham et al., 2016). The insights revealed from our data align with decades of work on gill specialization and ionoregulatory physiology, informing both experimental design and ecophysiological interpretation. We anticipate that this framework will facilitate comparative and stress-response studies in decapods and other aquatic invertebrates, including targeted tests of osmotic plasticity, transporter activity, ADP sensitivity, and ETS partitioning under ecologically realistic challenges.

## ACKNOLEDGEMENTS

This study was partially funded by the European Union’s Horizon Europe research and innovation program through a Marie Sklodowska Curie Global Fellowship (HE-MSCA-2021-PF grant agreement number 101062468-”MitoRescue”) awarded to G.A. Rivera-Ingraham. The authors would like to thank Dr. Olivia Holland (U. Griffith) for graciously providing access to two additional oxygraphs.

## ANIMAL ETHICS

All animals were captured, manipulated and euthanized humanely and following the International Guiding Principles for Biomedical Research Involving Animals, issued by the Council for the International Organizations of Medical Sciences. All manipulations were carried out under the animal ethics approval certificate ENV1723AEC number issued by Griffith University.

